# A novel mitochondrial regulon for Ferroptosis during fungal pathogenesis

**DOI:** 10.1101/2023.05.17.541075

**Authors:** Qing Shen, Fan Yang, Naweed I. Naqvi

## Abstract

Ferroptosis remains an underexamined iron- and lipid peroxides-driven cell death modality despite its importance to several human- and plant-diseases and to immunity thereof. Here, we utilized a range of chemical biological tools, genetic mutants and biochemical analyses to gain insights into how the fungal pathogen *Magnaporthe oryzae* undergoes Ferroptosis strictly in the spore cells to successfully transit to infectious development. We reveal a functional dependency between such intrinsic Ferroptosis and autophagy-mediated mitochondrial degradation. Mechanistically, the requirement of mitophagy for ferroptotic cell death was attributed to its ability to maintain a pool of metabolically active mitochondria. Pharmacological disruption of the electron transport chain or membrane potential led to complete inhibition of ferroptosis, thus simulating the loss of mitophagy phenotypes. Conversely, increased mitochondrial membrane potential in a mitophagy-defective mutant alleviated the Ferroptosis defects therein. Graded inhibition of mitochondrial Coenzyme Q (CoQ) biosynthesis with or without ferroptosis inhibitor Liproxstation-1 distinguished its antioxidant function in such regulated cell death. Membrane potential-dependent regulation of ATP synthesis and iron homeostasis, as well as dynamics of Aconitase in the Tricarboxylic acid cycle in the presence or absence of mitophagy, mitochondrial poisoning or iron chelation further linked mitochondrial metabolism to Ferroptosis. Lastly, we present an important mitochondrial bioenergetics and redox-regulatory network regulon essential for intrinsic Ferroptosis and its precise role in fungal pathogenesis leading up to the establishment of the devastating rice blast disease.

## Introduction

Ferroptosis is an evolutionarily conserved cell demise cause by iron-dependent peroxidation of membrane lipids that contain polyunsaturated fatty acid tail(s)^1-4^. Such nonapoptotic cell death was first reported in and/or well-studied in cancer cells, or engineered mammalian cell lines, and can be induced in several types of tumor cells through depletion of cellular glutathione (GSH) or inhibition of GSH-dependent glutathione peroxidase 4 (Gpx4) that directly converts the lipid hydroperoxides into nontoxic lipid alcohols^5,6^. A small molecule, Erastin, causes GSH depletion and Ferroptosis induction by inhibiting cellular uptake of cystine, which upon conversion to cysteine *in vivo* is used to synthesize GSH^6^. In tumor cells that are insensitive to Gpx4 inhibition, a second Ferroptosis resistance factor i.e. apoptosis inducing factor mitochondria-associated 2 (AIFM2) was identified and subsequently renamed as Ferroptosis suppressor protein 1 (FSP1)^7,8^. FSP1 is post-translationally modified and recruited to the plasma membrane where it reduces Coenzyme Q^7,8^ and vitamin K^9^ so as to stop phospholipid peroxidation via their radical-trapping antioxidant activities. STARD7 was subsequently identified as the CoQ transporter that shuttles between mitochondria and cytosol^10^, and thus unmasks the source of CoQ located near the plasma membrane.

Ever since the term Ferroptosis was coined, punctate/smaller mitochondria with increased membrane density were observed using transmission electron microscopy as the lone morphological marker^3^, but it remained enigmatic as what such morphological change means and what is the fate of such abnormal mitochondria. Ferroptosis can be induced by Erastin or the Gpx4 inhibitor RSL3 in 143B osteosarcoma cells devoid of mitochondria DNA^3^, thus questioning the importance of respiration-based reactive oxygen species (ROS) as the primary source of lipid peroxidation. Given the significant involvement of mitochondria in cellular iron and lipid metabolism^11,12^, it became important to investigate the mitochondria-Ferroptosis connection from these two perspectives.

Conversely, TCA and ETC have been found to enable Ferroptosis in mouse embryonic fibroblasts but only when it is specifically induced by cysteine starvation and not by Gpx4 inhibition^13^. It remains unclear as to why context-dependent requirement of intact mitochondria for Ferroptosis is evident in different cell types. One possibility could be that tumor cells, and probably also other cultured cell lines, modify cell metabolism in different ways for survival. For example, 143B osteosarcoma cells without mitochondrial DNA can rely on glycolysis for survival as far as the F1-ATPase capable of hydrolyzing ATP and the reversible ADP/ATP translocase named adenine nucleotide translocator are still functional^14^. Since membrane potential is necessary for import of most precursor proteins into the mitochondria ^15^ to enable proper metabolism therein, such ρ^0^ tumor cells also maintain sufficient MMP to sustain cell growth^14^. Thus, a clear and comprehensive understanding of the role of mitochondria in Ferroptosis also demands efforts in understanding the regulation of intrinsic Ferroptosis in healthy cells.

Rice plants, for example, undergo highly compartmentalized ferroptosis^17^, in epidermal cells invaded by the incompatible isolates of the blast fungus. Such strong immune response blocks and then kills the invading fungus within these dead cells. Interestingly, Ferroptosis also occurs in the spore/conidium of the rice blast fungus *M. oryzae* that triggers such host immune response, and represents a crucial determinant of the essential pathogenic development and thus the infection ability leading to the destructive blast disease^2,18^. Particularly, such intrinsic Ferroptosis initiates sequentially in the terminal conidial cell followed by the middle and proximal conidial cells (Fig. 1A)^18^, though the three cells in the conidium share the cytoplasm. Except gradient iron accumulation in the three cells^18^, little is known about the regulators controlling such precise Ferroptosis relay in the healthy interconnected cells.

**Fig. 1.**
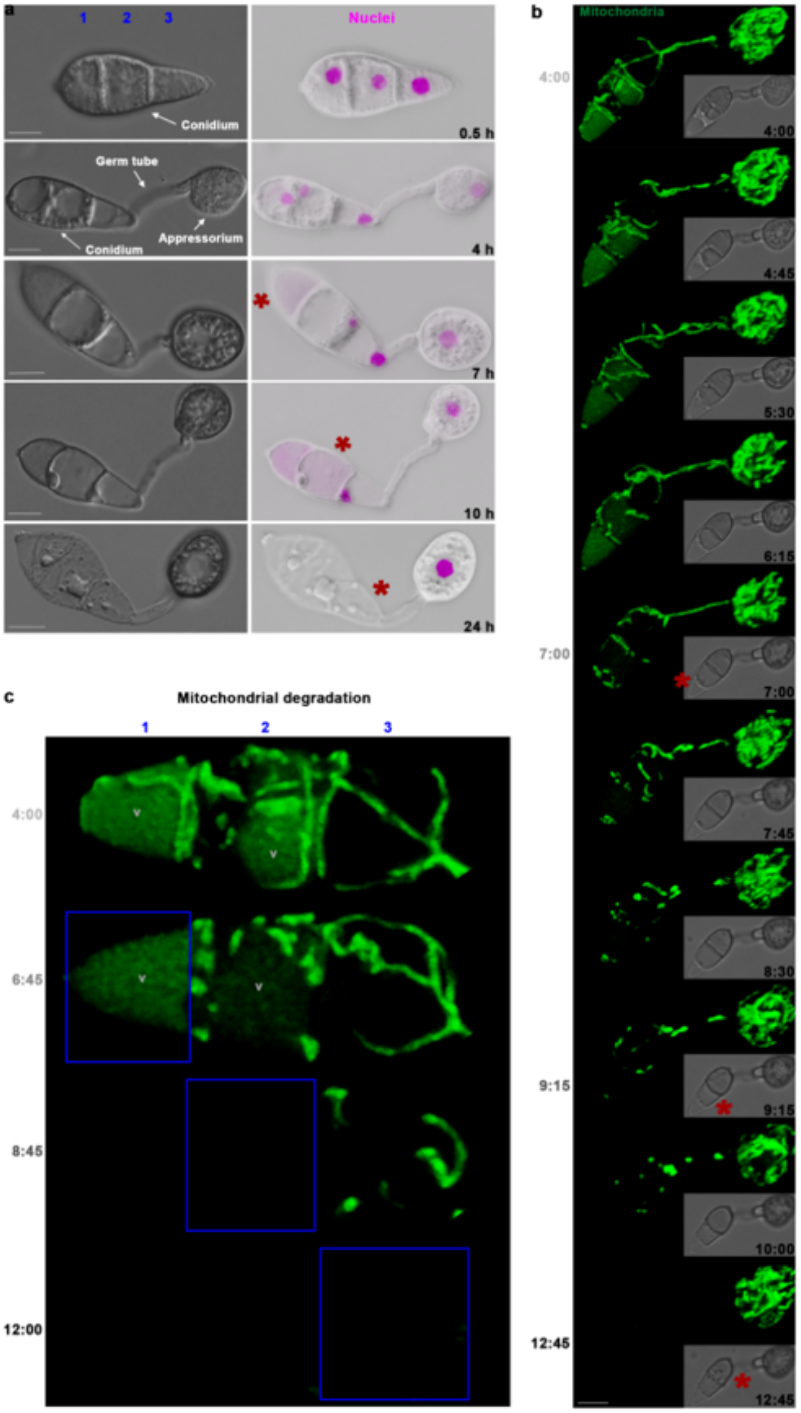
Regulated mitochondrial fission and degradation precede Ferroptosis in *M. oryzae* conidial cells. **a**, Programmed Ferroptosis occurs sequentially in the 3 connected conidial cells during the pathogenic development. Cells marked 1,2,3 in (**a**) and (**c**) are terminal, middle, and proximal conidial cells that initiate Ferroptosis first, second, and last, respectively, and time depicted as hours post inoculation (hpi). *M. oryzae* from 4 h to 24 h is shown in a way that from left to right are the three-celled conidium, germ tube, and appressorium. Red asterisk marks the dead cell, which is also indicated by the disappearance of the nucleus (hH1-GFP, color inverted) in the projection image. DIC images are single plane, and the scale bar equals 5 μm for both (**a**) and (**b**). The same experiment has been repeated 3 times with consistent results. **b**, Mitochondrial fusion, fission, and degradation along with sequential Ferroptosis events. Bright field images are single plane and overlapped in the 3D images of mitochondria displayed by a mitochondrial targeting sequence fused with GFP (MTS-GFP). Red asterisks mark collapsed conidial cells. Data shown is representative of 4 replicates of the experiment. **c**, Sequential mitochondrial degradation in the conidium correlates with Ferroptosis therein. Mitochondrial degradation finished first in the terminal cell (6:45 hpi), and then in the middle cell (8:45 hpi) and lastly in the proximal cell (12:00 hpi), precisely occurring before the sequential Ferroptosis cycle. Blue rectangles highlight the cells that just completed mitochondrial degradation. V: vacuole. Images in (**b**) and (**c**) are from the same time lapse experiment.

Iron that drives Ferroptosis in mammalian cells is found to be released by autophagic degradation of ferritin^19,20^, which is an iron storage protein complex. However, ferritin is not conserved in Magnaporthe albeit Ferroptosis therein being autophagy dependent^21^. Thus, whether other types of selective autophagy, such as mitophagy that targets dysfunctional mitochondria, bridge the gap between Ferroptosis and autophagy in fungi remain to be addressed. Here, using time lapse live-cell imaging, we report that mitochondrial fission and then vacuolar degradation occur prior to Ferroptosis initiation in the conidium of *M. oryzae* during pathogenic development. Mitophagy, which mediates such mitochondrial degradation, is shown to be essential for Ferroptosis and plays a crucial role in maintaining metabolically active mitochondria. Systematic investigation of the contributions of mitochondrial metabolism to Ferroptosis including Coenzyme Q and ATP synthesis, iron homeostasis and fatty acid catabolism provide a comprehensive view of a novel mitochondrial regulon essential for intrinsic Ferroptosis and its role in fungal pathogenesis that enables the establishment of the devastating rice blast disease. Such knowledge highlights the conserved Ferroptosis as a novel target for crop protection, and in turn will help better understand the mitochondria-Ferroptosis nexus in human diseases too.

## Results

### Mitochondrial fission and degradation precede intrinsic Ferroptosis cell death in *M. oryzae*

Pathogenesis in *M. oryzae* is a spatio-temporally controlled developmental process featuring the formation and maturation of the infection structure called the appressorium (Fig. 1a). As a critical and programmed step of such development, the 3 connected cells in the asexual spore or conidium undergo Ferroptosis individually and sequentially (Fig. 1a). Interestingly, a similar sequential behavior prior to ferroptotic cell death was observed for mitochondrial degradation, which commenced and completed first in the terminal cell prior to its collapse and death, and then the same two processes repeated subsequently in the middle and proximal cells around 2 and 5 hours later, respectively (Fig. 1b, c and Extended Data Fig. 1a). In each of the dying conidial cells, mitochondria showed a trend changing from tubular filaments to punctate structures, thus suggesting the involvement of the fission process prior to selective degradation in the vacuole (Supplementary Movie 1). The occurrence of Ferroptosis specifically correlated with and followed such precise mitochondrial degradation thus underscoring the importance of this organellar homeostasis in regulated cell death early in the infection cycle.

### Mitochondrial degradation via mitophagy is necessary for ferroptotic cell death

Mitophagy, which targets dysfunctional or excess mitochondria for vacuolar degradation using the autophagy machinery, is one of the pathways responsible for such organellar turnover and homeostasis^22^. Weak free GFP signal in the vacuole (Fig. 1c) implied that mitophagy could be an important pathway responsible for the observed mitochondrial degradation. Therefore, we examined Ferroptosis in the mitophagy-deficient mutant, *atg24*Δ^22^, and found that loss of mitophagy (Fig. 2a) significantly suppressed Ferroptosis and consequently increased the conidial viability in *atg24*Δ conidia as compared to the wild-type *M. oryzae* (Fig. 2b). An extensively fused and dense network of mitochondria was present in the mitophagy-defective *atg24*Δ conidial cells but was rarely seen in the wild-type Magnaporthe conidia wherein the mitophagy was fully functional (Fig. 2a). Next, mitophagy was disrupted by pharmacologically inhibiting mitochondrial fission thus resulting in stably fused mitochondria that persist and are unable to be cleared via mitophagy due to the larger size and tubular constraints. As expected, disrupting mitochondrial fission led to similar Ferroptosis inhibition in a dose-dependent manner (Fig. 2c). Likewise, lipid peroxides which drive Ferroptosis failed to accumulate along the outer membranes in the *atg24*Δ conidia (Fig. 2d) and rendered the *atg24*Δ conidia (that consequently lack Ferroptosis) incapable of causing blast infection in rice plants (Fig. 2e) as compared to the wild-type *M. oryzae* isolate. Together, these results confirmed the functional correlation between Ferroptosis and the precise vacuolar degradation of mitochondria, thus highlighting an essential role for mitophagy therein.

**Fig. 2.**
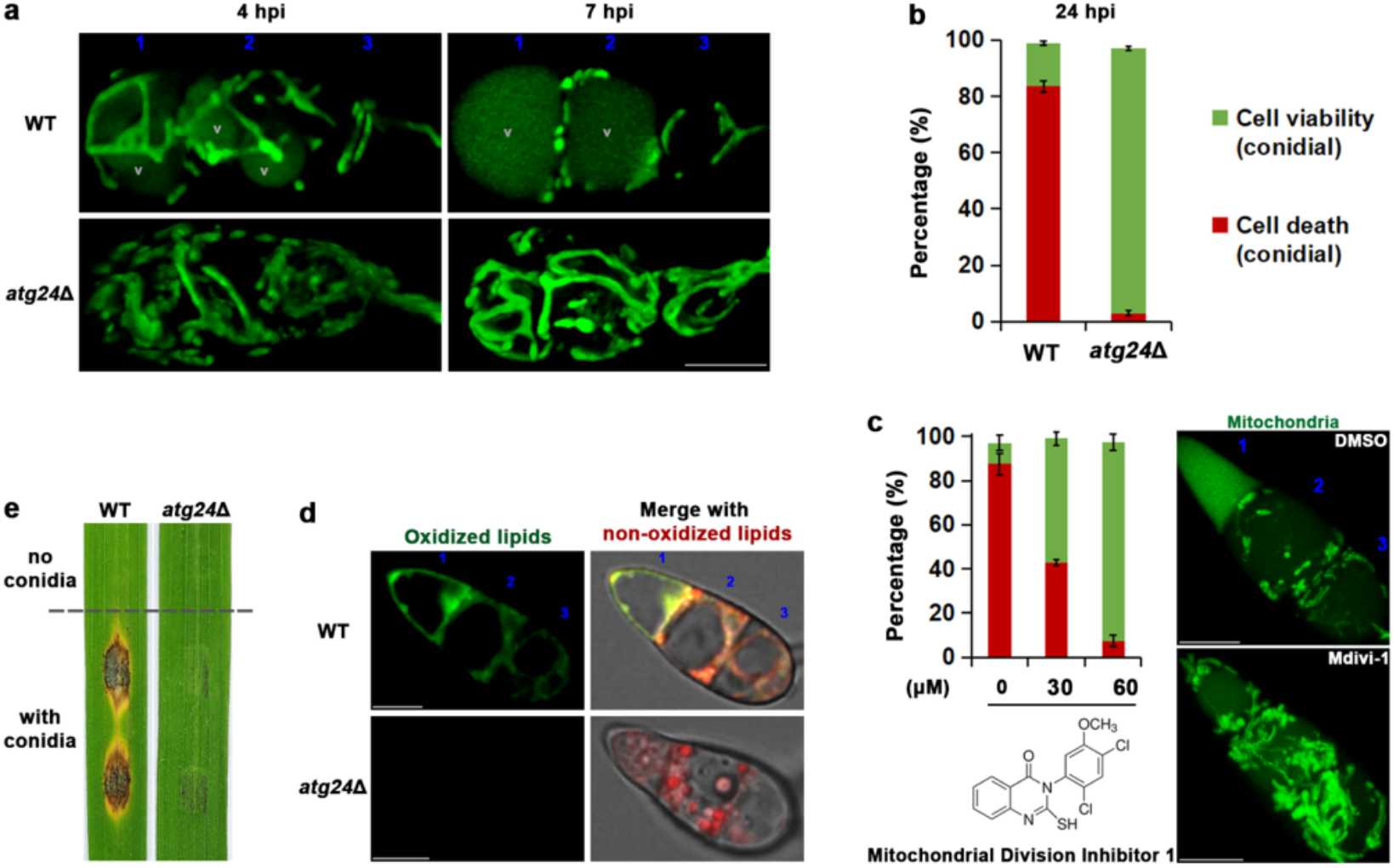
Disruption of mitochondrial degradation via mitophagy suppresses ferroptotic cell death. **a**, Mitochondria fuse instead of undergoing degradation when mitophagy is defective. Mitochondria (MTS-GFP shown as 3D images) in wild type (WT) or mitophagy mutant *atg24*Δ were observed at 4 or 7 hpi. V for vacuole. **b**, Ferroptosis fails to occur in *atg24*Δ conidia. Conidial cell death or viability was quantified at 24 hpi and depicted as mean ± *SD* from 3 technical replicates, each contains 100 conidia for both wild type (WT) and *atg24*Δ. **c**, Chemical disruption of mitophagy through Mitochondrial division inhibitor-1 (Mdivi-1) also suppresses Ferroptosis in a dose-dependent manner. Effectiveness of Mdivi-1 (60 μM) on mitochondrial fission was verified via MTS-GFP (projection) at 7 hpi. Conidial cell viability (green) or death (red) was quantified at 24 hpi and displayed as mean ± *SD* (3 technical repeats, n=100 each per dose). DMSO (0.1%): solvent control. **d**,**e**, Mitophagy defective *atg24*Δ conidia fail to accumulate lipid peroxides (oxidized lipids) to a level that can cause Ferroptosis and are defective in Ferroptosis-dependent blast disease infection in rice. **d**, Oxidized (green) and non-oxidized (red) variants of lipids in wild-type (WT) or *atg24*Δ conidial cells were observed at 7 to 8 hpi via C11-BODIPY^581/591^ staining and ratiometric epifluorescence confocal microscopy and shown as single plane images. **e**, Rice blast lesions photographed at 7 days post inoculation. Data shown from (**a**) to (**e**) are representative of 3 independent replicates of the experiment. Scale bar = 5 μm for (**a**), (**c**), and (**d**).

### Mitophagy maintains a pool of metabolically active mitochondria for Ferroptosis

To address the exact function of mitophagy during such developmental cell death, we compared the mitochondrial metabolism in wild type and *atg24*Δ using the fluorescent dye TMRE whose mitochondrial localization depends on membrane potential. Interestingly, although a robust filamentous mitochondrial network persisted in the *atg24*Δ conidia, such mitochondria showed a massive reduction in the membrane potential as compared to those in the wild-type *M. oryzae* (Fig. 3a), which indicated that mitophagy is required for maintaining the overall metabolic activity.

**Fig. 3.**
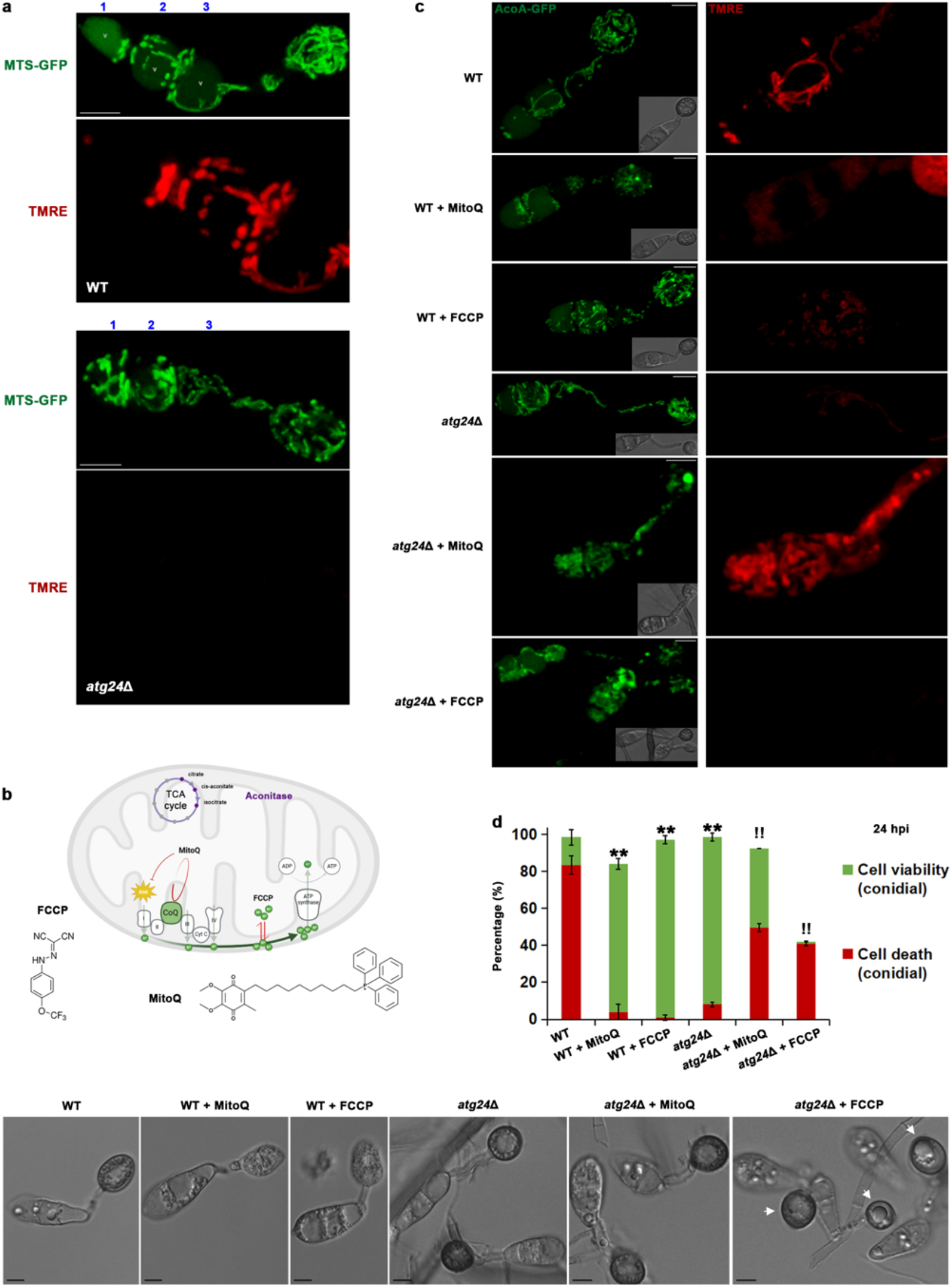
Mitophagy maintains a pool of metabolically active mitochondria that are important for Ferroptosis. **a**, Mitochondrial membrane potential as detected by TMRE reduces dramatically in the *atg24*Δ conidia. MTS-GFP outlines mitochondria at 6 hpi. Conidia of wild type (WT) or *atg24*Δ were examined in three independent experiments with at least 30 conidia in total for each strain. V, marks the vacuole. Images shown are all projections. **b**, Schematic model summarizing the mode of action for MitoQ, FCCP, and Aconitase. CoQ, ubiquinone/Coenzyme Q. ROS, reactive oxygen species. The graphic was created using BioRender. **c, d**, Positive correlation between mitochondrial metabolism and Ferroptosis. **c**, Mitochondria in wild type or *atg24*Δ (*AcoA-GFP atg24*Δ #36) are displayed by AcoA-GFP, and TMRE reflects the mitochondrial membrane potential. Images for both panels are projections, whereas the inserted DIC images are single plane. **d**, Conidial cell viability or death was quantified at 24 hpi and displayed as mean ± *SD* from 3 technical replicates with n=100 conidia for each treatment per strain per replicate. ** and !! means *P*<0.01 as compared with WT and *atg24*Δ, respectively. Single plane DIC images showing the morphologies of wild type or *atg24*Δ with or without MitoQ or FCCP at 24 hpi. White arrows mark abnormal appressoria. Data in (**c**) and (**d**) are representative of 2 independent replications showing consistent results. Total 600 conidia were quantified and at least 26 conidia were examined per treatment per strain for quantification and imaging, respectively. Scale bar equals to 5 μm for (**a**), (**c**) and (**d**).

Indeed, time lapse imaging showed that all filamentous and punctate mitochondria in the mitophagy-competent wild-type strain, are metabolically active and stain positive with TMRE (Extended Data Fig. 1b). Consistent with this observation, localization of the mitochondrial matrix protein Atp1, the α subunit of the F1 fraction of the ATP synthase, further confirmed that mitophagy helps keep and maintain a pool of metabolically active mitochondria prior to the initiation of ferroptotic cell death (Extended Data Fig. 1c). Together, these data verified the role of mitophagy in maintaining active mitochondria and pointed out the necessity to understand the relationship between mitochondrial metabolism and Ferroptosis.

Therefore, the electron transport chain was disrupted by replacing ubiquinone/CoQ with its analog Mitoquinone (MitoQ) or the membrane potential was directly decreased through the protonophore FCCP (Fig. 3b). A substantial decrease in Ferroptosis along with a dramatic decline in membrane potential was evident upon such disruptive treatments (Fig. 3c, d). Such outcomes were reminiscent of and akin to the phenotypic defects associated with the loss of mitophagy in *M. oryzae* (Fig. 3c, d).

CoQ is synthesized in mitochondria and exported to the plasma membrane where it inhibits Ferroptosis as an antioxidant ^7,8,10^. Thus, the role of CoQ as an electron carrier in regulating Ferroptosis was further verified using another analog Idebenone and the inhibitor of the ETC Complex III, Antimycin A, was included for comparison (Fig. 4a). Both drugs suppressed conidial cell death to a similar level (Fig. 4b). Furthermore, inhibiting CoQ biosynthesis via 4-CBA using a higher dose also suppressed Ferroptosis (Fig. 4c). However, lower doses of 4-CBA, probably by reducing the pool of CoQ near the plasma membrane, promoted Ferroptosis which could be reversed by the antioxidant Liproxstatin-1 (Fig. 4c, d). The function of CoQ as a potent antioxidant implies that it may be able to protect the mitophagy mutant from the sudden loss of membrane potential if such a decrease was caused by oxidative stress or damage and such impairment remained salvageable. Indeed, treatment of *atg24*Δ with MitoQ significantly increased its MMP and alleviated the Ferroptosis defect as judged by the increase in the percentage of *atg24*Δ conidia that showed wild-type-like features i.e. conidial cells dead but appressoria viable and mature (Fig. 3c, d). As a negative control, FCCP had no obvious effect on *atg24*Δ MMP and failed to rescue the Ferroptosis defect of *atg24*Δ (Fig. 3c, d). Instead, it triggered a non-ferroptotic cell death accompanied by an abnormal spread of cell death to the appressoria (Fig. 3d).

**Fig. 4.**
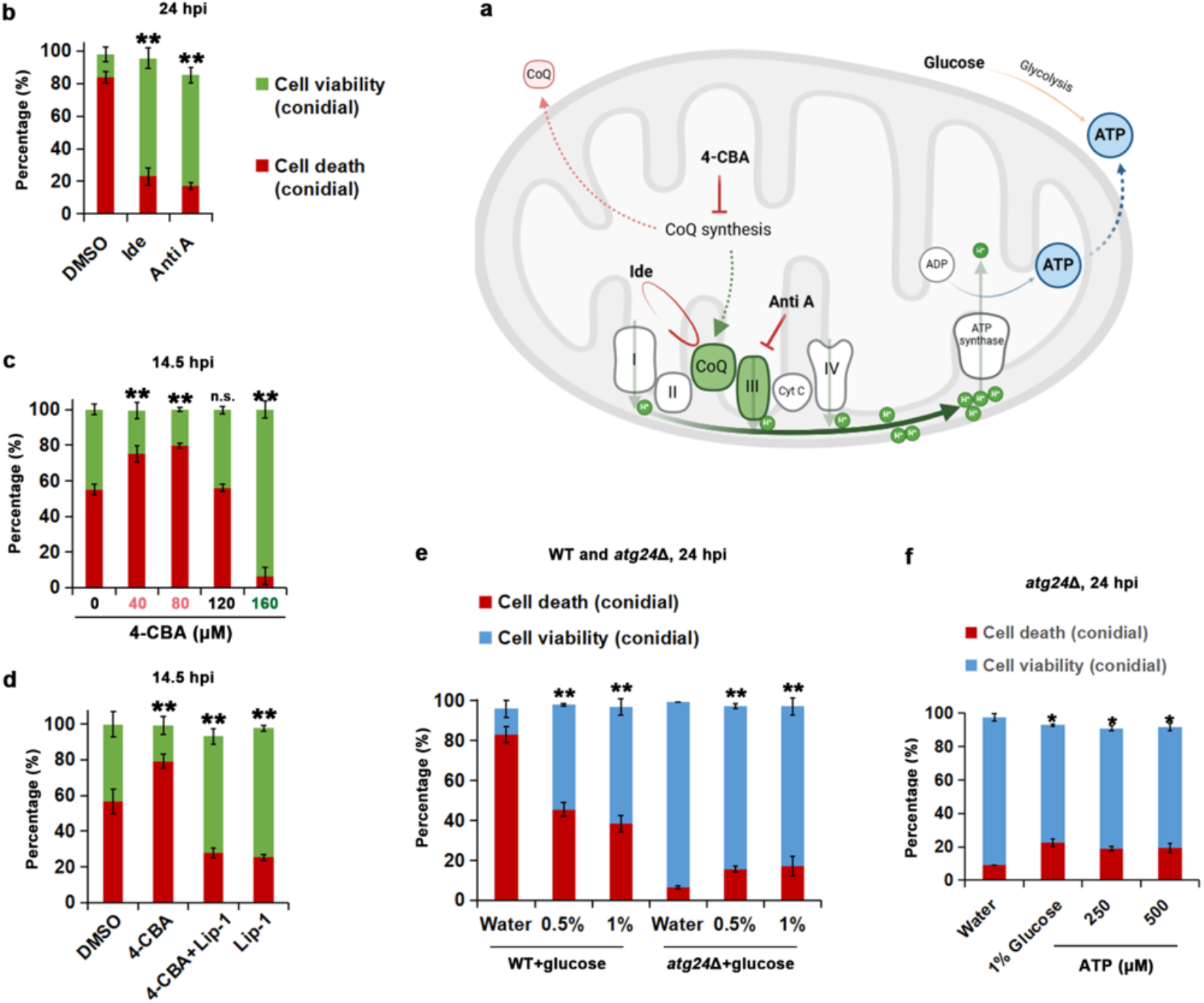
Coenzyme Q and ATP modulate Ferroptosis. **a**, Model summarizing the relationship between mitochondrial metabolism and fungal Ferroptosis. **b**, Disruption of electron transport chain leads to Ferroptosis suppression. Conidial cell viability/death was quantified at 24 hpi and is displayed as mean ± *SD* from 3 technical replicates with n=100 conidia for each treatment per replicate. **c**, Inhibition of Coenzyme Q biosynthesis via exogenous 4-CBA promotes Ferroptosis at lower doses, whereas suppresses it at a higher dose. Conidial cell viability (green) or cell death (red) was quantified at 14.5 hpi and is displayed as mean ± *SD* derived from 3 technical replicates (n=100 conidia each) of the experiment for each dose. **d**, Ferroptosis induction caused by 4-CBA is reversed by Liproxstation-1 (Lip-1). Conidial cell viability/death presented as mean *± SD* (3 technical replicates, n=100 conidia for each treatment per replicate) was quantified at 14.5 hpi. **e**, Effect of glucose on conidial viability in the wild-type (WT) or *atg24*Δ strain of *M. oryzae*. Death (red) or viability (blue) of the conidium was quantified at 24 hpi and presented as mean ± *SD* derived from 3 technical replicates, each containing 100 conidia per treatment per strain. **f**, Ferroptosis defect in *atg24*Δ is slightly suppressed by increasing glycolysis (via glucose) or via exogenous provision of ATP. Conidial cell death (red) or viability (blue) in *atg24*Δ was quantified at 24 hpi and presented as mean ± *SD* (3 technical replicates, n=100 conidia for each treatment per replicate). For quantification in (**b**) to (**f**), ** (p < 0.01) and * (p < 0.05) indicate significant differences, while n.s. refers to not significant as compared to the solvent (DMSO or water) control, and data presented have been confirmed through 2 or 3 biological repeats of the experiments.

To verify and further understand the relationship between mitochondrial metabolism and Ferroptosis, a native in locus epifluorescence-tagged strain of the mitochondrial Aconitase (named Aconitase A in *M. oryzae*; AcoA for short) which catalyzes the transition of citrate to isocitrate via cis-aconitate in the TCA cycle was made (Fig. 3b and Extended Data Fig. 2a-c), which behaved just like the wild type in terms of the ability to infect rice plants (Extended Data Fig. 2d). Loss of *ATG24*, significantly affected the mitochondrial localization of AcoA (Extended Data Fig. 2e-g), suggestive of defects in MMP-associated import of freshly synthesized AcoA into mitochondria. In line with this finding, AcoA marked mitochondria became fragmented with MitoQ or FCCP treatment (Fig. 3c), supporting the disruption of mitochondrial metabolism, most likely also indirectly affects the TCA cycle, with these mitochondrial poisons, which again is consistent with the decrease in membrane potential observed with such drug treatments (Fig. 3c). Taken together, our data revealed a requirement for mitochondrial membrane potential-dependent metabolism in proper and timely induction of Ferroptosis.

### Cellular ATP and iron link mitochondrial metabolism to Ferroptosis

Next, we asked whether mitochondrial metabolism and Ferroptosis are functionally linked. ATP synthesis was first tested by promoting glycolysis via exogenous glucose in the low MMP *atg24*Δ or by directly providing ATP *in trans*. Interestingly, increase in the cellular ATP levels slightly but significantly increased the conidial cell death in the *atg24*Δ regardless of the dosage of glucose or ATP (Fig. 4e, f), which further underscored the role of ATP synthesis and mitochondrial metabolism in Ferroptosis.

An overall iron deficiency was always evident in the *atg24*Δ mutant (Fig. 5a), which further explains its consequent defect in iron dependent Ferroptosis. Short term iron chelation through CPX caused an increase in MMP in the wild-type Magnaporthe, and such mitochondrial response to iron shortage was also observed in *atg24*Δ although to a much lower degree (Fig. 5b), which implies a role of active mitochondria in mediating iron homeostasis and may account for the iron deficiency observed in the *atg24*Δ (Fig. 5a). AcoA requires iron as a cofactor and is sensitive to iron deficiency. Indeed, iron chelation caused mild fusion in AcoA-GFP mitochondria in the wild type and showed strong fragmentation of mitochondria in *atg24*Δ (Fig. 5b), thus reflecting the different degrees of iron deficiency sensed by AcoA. Intriguingly, iron deficiency in the *atg24*Δ, unlike the one caused by iron chelation/CPX treatment, could not be alleviated by direct iron supplementation as the *atg24*Δ failed to undergo Ferroptosis with or without such iron supplementation (Fig. 5c), thus reflecting a unique iron starvation phenotype of such low MMP in *atg24*Δ.

**Fig. 5.**
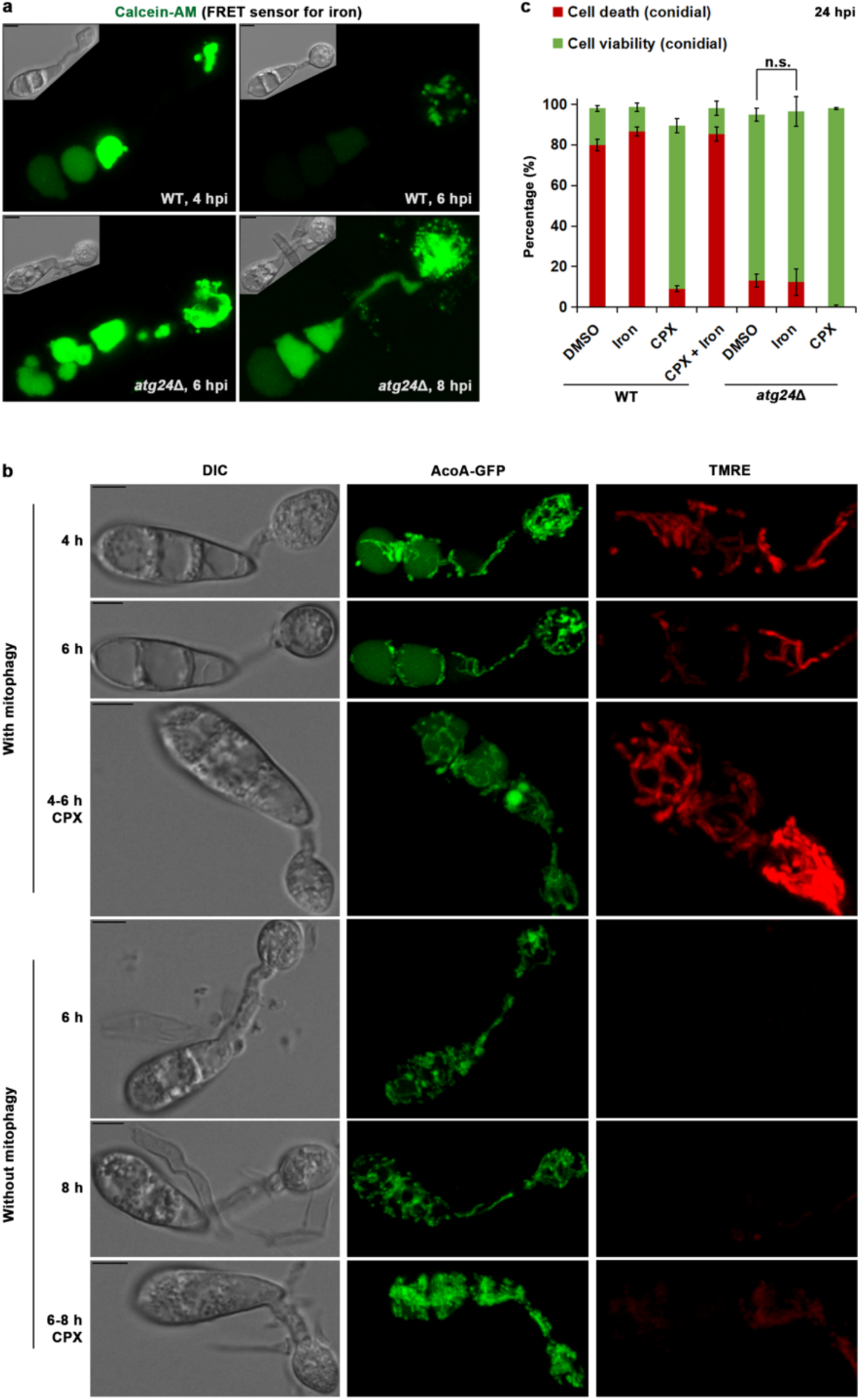
A role of active mitochondria in mediating cellular iron homeostasis. **a**, Iron deficiency in the *atg24*Δ as shown by the FRET sensor Calcein-AM as projection images at indicated time points. Insets are projection of DIC images. WT: wild type. **b**, Change of mitochondrial membrane potential and dynamics of AcoA in response to iron starvation in the presence and absence of mitophagy. Mitochondrial membrane potential in wild-type conidia reflected by TMRE increases when iron is cheated by CPX, and such change becomes very small when mitophagy is lost. Projection images are shown except DIC, which is single plane. *AcoA-GFP#3* and *AcoA-GFP atg24*Δ #36 were used to show the mitochondrial changes. **c**, Ferroptosis defect of *atg24*Δ cannot be suppressed by exogenous ferric ion (FeCl3). Conidial cell death/viability of wild type (WT) or *atg24*Δ in the presence or absence of iron and/or the iron chelator CPX was quantified at 24 hpi and is depicted as mean ± *SD*, which is derived from 3 technical replicates, each containing 100 conidia per treatment per strain, and n.s. refers to no significant difference. Calcein-AM staining (**a**) was repeated thrice, mitochondrial response to iron chelation (**b)** was repeated twice, and the experiments for *atg24*Δ conidial cell death with or without iron or CPX (**c**) were repeated at least 5 times. Scale bar equals 5 μm for images in (**a**) and (**b**).

### Mitochondrial β-oxidation is not essential for Ferroptosis

Catabolism of PUFA in the mitochondria was also examined in *M. oryzae* using the mitochondrial β-oxidation mutant *ech1*Δ^23^, which shows defects in fatty acid oxidative catabolism^23^. Surprisingly, such mutant was capable of undergoing Ferroptosis just like the wild-type *M. oryzae* (Fig. 6a).

**Fig. 6.**
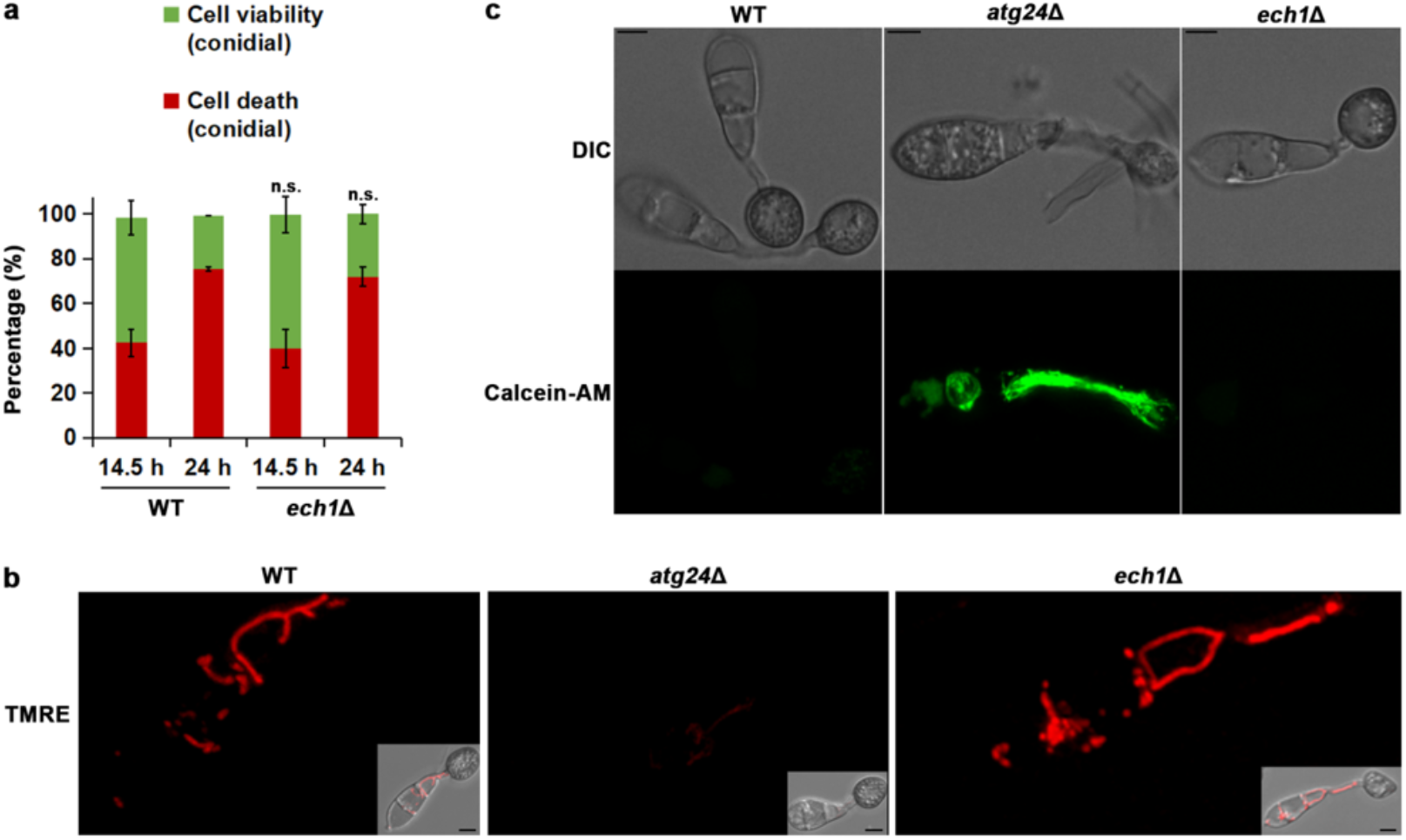
The mitochondrial β-oxidation mutant *ech1*Δ is not defective in Ferroptosis. **a**, Conidial cell viability or death in wild type (WT) or *ech1*Δ was quantified at the indicated time points and presented as mean ± *SD* (3 technical replicates, n=100 conidia for each time point per strain per replicate). n.s. means no significant difference was detected as compared to the WT at the corresponding time points. Experiment has been repeated twice. **b**, Mitochondrial metabolism reflected by mitochondrial membrane potential (TMRE/projection, only 3 conidial cells are shown) is dramatically reduced in *atg24*Δ, but remains normal in *ech1*Δ, as compared to the wild-type *M. oryzae* (WT). Insets are full view projections merging bright field and TMRE images. **c**, Cellular iron availability shown by Calcein-AM at 7 hpi indicates that *ech1*Δ, unlike *atg24*Δ, does not suffer from iron deficiency as compared to the wild type (WT) Magnaporthe. DIC images are single plane, while Calcein-AM images are projections. Scale bars = 5 μm for (**b**) and (**c**).

Furthermore, no change in mitochondrial membrane potential or cellular iron availability was observed in the *ech1*Δ mutant as compared to the wild type (Fig. 6b, c), which in turn supports the membrane potential-dependent roles of mitochondria in cellular iron control and Ferroptosis. In conclusion, results here demonstrate that mitophagy helps maintain a pool of metabolically active mitochondria that are important for Ferroptosis and unveil an important mitochondrial bioenergetics and redox regulatory network for precise ferroptotic cell death in the rice blast pathosystem.

## Discussion

The process of mitophagy likely distinguishes between functional and dysfunctional mitochondria based on low levels of mitochondrial membrane potential and clears such damaged organelles via vacuolar degradation in a precise spatio-temporal manner in Magnaporthe. This is important for ensuring and enabling a pool of metabolically active mitochondria in the viable conidial cells during the early stages of infection-related development. Such active mitochondria are, in turn, necessary for the programmed Ferroptosis during pathogenesis, with the membrane potential-dependent modulation of cellular iron homeostasis and/or bioavailability and/or ATP synthesis as promising links connecting these two important processes.

Increase in mitochondrial membrane potential has also been observed in mouse embryonic fibroblasts undergoing Ferroptosis and is attributed to cysteine deprivation therein^13^. It is worth noting that cysteine is also required for the synthesis of iron-sulfur clusters in mitochondria in a membrane potential dependent manner^11^, thus linking the mitochondrial responses to iron shortage observed in Magnaporthe (Fig. 5b). Proper function of the TCA cycle and the electron transport chain, likewise, depends on iron as a cofactor, which intricately links such iron starvation responses and explains the dependence of Ferroptosis on iron from a new perspective given the requirement of mitochondrial metabolism and membrane potential in mounting such a specific regulated cell death modality.

Unlike mitochondria in mitoQ-or FCCP-treated wild type, extensively fused mitochondria were also observed in the *atg24*Δ mutant, thus suggesting that mitochondrial fusion is likely a salvage option to titrate and reduce the negative effects exerted by damaged or dysfunctional mitochondria with reduced membrane potential that are likely substrates for mitophagy. Supporting such a hypothesis, mitochondrial fusion was recently shown to restore the mitochondrial defects caused by mitochondrial DNA mutations and oxidative stress^24^. Interestingly, Ferroptosis was also found to be suppressed by such drug-induced mitochondrial fusion^24^, most likely by disrupting the timely clearance of dysfunctional mitochondria through mitophagy. The finding that mitoQ can alleviate *atg24*Δ defects is interesting and suggests that *atg24*Δ suffers similarly from oxidative stresses.

Thus, the inability of *atg24*Δ to execute Ferroptosis properly implies that mitochondrial ROS is unlikely to be a pro-Ferroptosis factor in *M. oryzae*. Overall, our data highlight the mitochondrial membrane potential-enabled metabolism as an important regulator of the developmental ferroptotic cell death in the rice blast fungus and provides a likely strategy for intervention of the devastating blast disease in cereal crops.

## Methods

### Fungal strains and growth conditions

*Magnaporthe oryzae* strains were cultivated on Prune Agar (PA) medium for vegetative growth and conidiation as described^25^. Blast isolate B157 obtained from the Directorate of Rice Research (Hyderabad, India) was used as the wild-type strain of choice. Epifluorescence-tagged strains including Histone H1 (hH1)-GFP^18^, MTS-GFP^23^, Atp1-GFP^23^, as well as deletion mutants *atg24*Δ with or without MTS-GFP^22^, and *ech1*Δ^23^ have been described in our previous publications.

### Generation of *AcoA-GFP* and *AcoA-GFP atg24*Δ strains

The plasmid construct for generating *ACOA-GFP* in locus tagged strain (Extended Data Fig. 2a) was made using the Clon Express MultiS One Step Cloning Kit (Vazyme, C113). Briefly, 4 overlapping fragments covering a part of *ACO A* (MGG_03521) exon3 (1132 bp just before the stop codon), a short sequence encoding a 4 amino acid linker, eGFP encoding sequence without start codon, Basta resistance cassette in the reverse direction, and a part of *ACO A* 3’UTR (1258 bp including the *ACO A* stop codon) was PCR amplified using primers listed in supplementary table 1 (primer 1 to 8) and purified. Meanwhile, the empty vector pFGL815 (Addgene Plasmid #52322) was linearized using Xma1 and Xba1, and subsequently used to assemble the 4 PCR fragments using the MultiS One Step Cloning Kit. Same technology, empty vector, and kit were used for generating the *ATG24* deletion construct as indicated in Extended Data Fig. 2e using primers listed in supplementary table 1. The 2 resultant plasmids were sequence verified and then used to transform AGL1 Agrobacterium individually through electroporation. Wild type B157 was the parent strain for generating the *ACOA-GFP* strain, while *ACOA-GFP* #3 was selected for subsequent *ATG24* deletion. Agrobacterium T-DNA mediated transformants of the rice-blast fungus *M. oryzae* were selected on basal medium (BM) with Basta (Glufosinate-ammonium, Dr. Ehrenstorfer GmbH) or sulfonylurea (Chlorimuron ethyl, Dr. Ehrenstorfer GmbH)^25^. Positive transformants were further verified though epifluorescence imaging and PCR using primers indicated in Extended Data Fig. 2a,e, which are also listed in supplementary table 1.

### Stains, and confocal microscopy

Sterile water droplets (20 μl) containing freshly harvested conidia at a concentration of 1.5×10^5^ conidia/ml were inoculated on hydrophobic cover glass (Menzel-Glaser or Matsunami) for normal imaging, or a 30 μl droplet at the concentration of 1×10^5^ conidia/ml were used for inoculating in glass-bottom culture dishes (MatTek Corporation, P35G-0-14-C) for time lapse imaging. Roughly 20-30 min before imaging, conidia were stained with 10 µM C11-BODIPY^581/591^ (Thermo Fisher, D3861) to detect lipid peroxidation, or with 250 nM Tetramethylrhodamine ethyl ester perchlorate (TMRE, Sigma, 87917) to assess mitochondrial membrane potential, or 1 µM Calcein-AM (C3099; Invitrogen) to test cellular iron availability.

Laser scanning confocal microscopy was performed using the Leica TCS SP8 X inverted microscope system (Leica Microsystems) under the control of Leica Application Suite X software package (release version 3.5.7.23225). Experiments with C11-BODIPY^581/591^ were done using Matsunami micro slide glass (Matsunami, S7213) and an HCX Plan Apochromat lambda blue 63×/1.20 water immersion objective. Argon laser (excitation, 488 nm; emission, 500-535 nm) was used for the oxidized form whereas the white light laser (excitation, 561 nm; emission, 573-613 nm) was used for non-oxidized variant. HC Plan Apochromat CS2 100× or 63×/1.4 oil immersion objectives and white light laser were used for GFP (excitation, 488 nm; emission, 500-550 nm), Calcein-AM (excitation, 494 nm; emission, 510-550 nm), and TMRE (excitation, 540 nm; emission, 580-610nm). All the lasers associated with Leica TCS SP8 were controlled by the AOTF (Acousto-Optical-Tunable-filter), and fluorescence images were captured using the Leica Hybrid Detector as Z stacks of 10 to 25 sections (0.5 µm-spaced). Time lapse images were further processed using the IMARIS v.9.6.0 software (Bitplane AG, Zurich, Switzerland).

### Pharmacological treatment and cell viability/cell death measurements

Conidia were inoculated on cover glass as above described. The 4-Chlorobenzoic acid (4-CBA) (Sigma, 135585) or glucose (Duchefa Biochemie, G0802) was added at 0 hour post inoculation (hpi), ATP (Sigma,A6419) was added at 2.5 hpi, while Mdivi-1 (Sigma, M0199), 4 µM Mitoquinone (MitoQ, Cayman chemical, 29317), 2 µM FCCP (Sigma, C2920), 32 µM Idebenone (Cayman chemical, 15475), 9.4 µM Antimycin A (Sigma, A8674), 54 µM liproxstatin-1(Sigma, SML1414), 5 µM ciclopirox olamine (Sigma, C0415), or 5 µM FeCl3 (Sigma, F2877) was added at 4 hpi to maximally restrict the chemical effect on Ferroptosis or conidial cell death and minimize the off-target effects on other processes. For the same reasons, MitoQ or FCCP was added to *atg24*Δ conidia at 7 hpi. Conidial cell viability or cell death was quantified using Trypan blue at the indicated time points. Conidia capable of appressorium formation and possessing 1 to 3 viable conidial cells were considered “viable”, while those having a viable appressorium but all the 3 conidial cells inviable were regarded as “dead”. Statistical analysis was achieved via Student’s *t*-test.

### Rice Blast infection assays

The youngest leaf of susceptible CO39 rice seedlings at 4 to 5 leaf stage was used for testing rice infection by wild type, *atg24*Δ or *AcoA-GFP* #3, and blast lesions examined at 7 days after inoculation as previously described^18^.

## Additional information

**Supplementary Movie 1**. Related to Fig. 1b and Extended Data Fig. 1a.

Time-lapse imaging of the dynamics and vacuolar degradation of mitochondria during conidial cell death in *Magnaporthe oryzae*. Precise mitochondrial fission and mitophagy precede ferroptotic cell death in each conidial cell in a sequential manner starting with the cell most distal to the appressorium. Mitochondria (MTS-GFP) were monitored from 4 to 12:45 hpi and imaged every 15 min. Merge of MTS-GFP and bright field (BF) is shown as a 3D rendition. The brightness of BF images was manually adjusted to demonstrate the collapse of the dead cells. Images in Fig 1b,c, Extended Data Fig. 1a, and here are from the same time lapse experiment.

**Supplementary Table 1**. Primers used for plasmid construction and strain verification.

## Acknowledgements

We thank the Fungal Patho-Biology Group (TLL, Singapore), Yanjun Kou (CNRRI, China) and Yizhen Deng (SCAU, China) for discussions and useful suggestions. This research was funded by intramural grants from the Temasek Life Sciences Laboratory, Singapore.

## Author contributions

Qing Shen: experimental investigation, data curation, methodology, validation, co-writing original draft manuscript. Fan Yang: strain development, validation, methodology, microscopy. Naweed I. Naqvi: conceptualization, formal analysis, funding acquisition, project administration, resources, supervision, co-writing original draft, reviewing and revisions.

## Conflict of interest

The authors declare that there is no conflict of interest involved.

---

**Extended Data Fig. 1.**
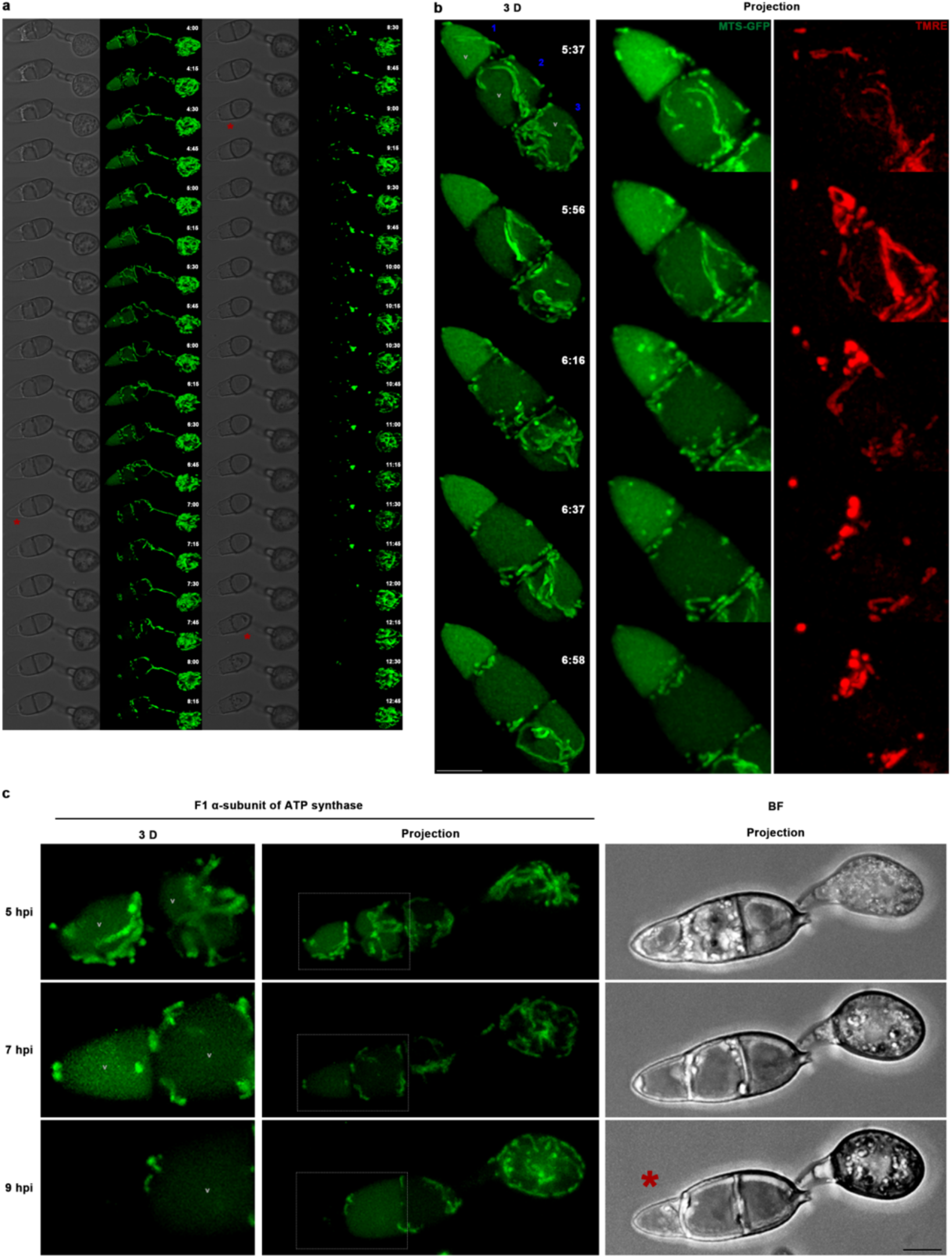
Timely mitochondrial clearance maintains a pool of active mitochondria. **a**, Montage showing fusion, fission, and degradation of mitochondria along with conidial Ferroptosis. Bright field image is single plane while MTS-GFP is shown as a rendered 3D. Collapsing or dead conidial cells are highlighted by red asterisks. **b**,**c**, When mitophagy functions properly, both punctate and filamentous mitochondria are metabolically active. Time lapse imaging was performed to assess mitochondrial membrane potential. MTS-GFP outlines mitochondria in (**b**). Gray rectangle marks the origin of the enlarged terminal and middle cells, and the red asterisk indicates the collapsed terminal cell in (**c**). V denotes the vacuole. BF stands for bright field. Experiments in (**b**) and (**c**) were repeated thrice independently with 6 conidia in each instance. Scale bars, 5 μm.

**Extended Data Fig. 2.**
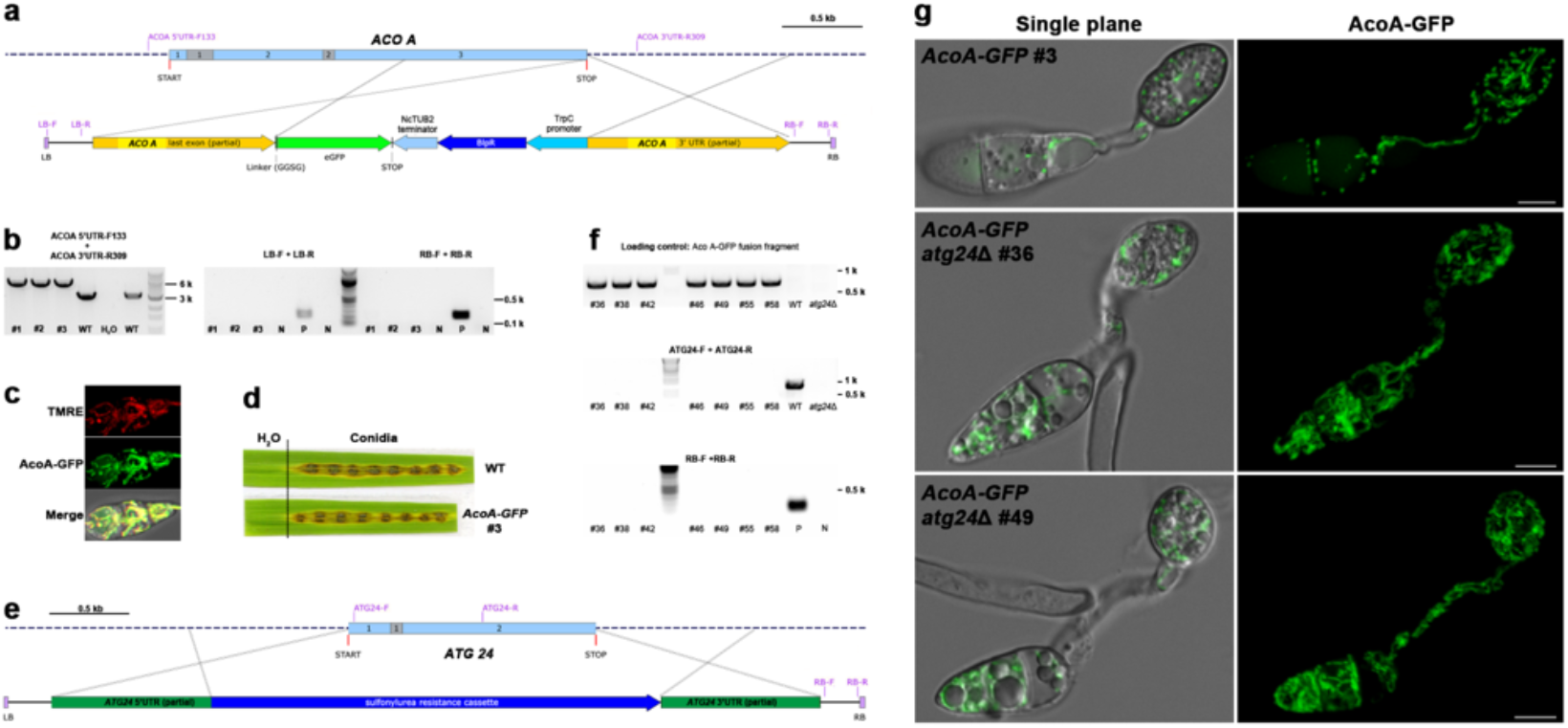
In locus GFP tagging of the mitochondrial Aconitase and subsequent *ATG24* deletion. **a**,**b**,**e**,**f**, In locus GFP tagging of the mitochondrial Aconitase and subsequent *ATG24* deletion. **a**, Insertion of sequences expressing GFP and Bialaphos resistance (Bar) cassette, which is driven by *Aspergillus nidulans TRPC* promoter and terminated with *Neurospora crassa TUB2* terminator, through homologous recombination at the *ACONITASE A* (*ACO A*) locus just before its stop codon. ACOA 5’ UTR-F133 and ACOA 3’ UTR-R309 are primers used to verify in locus insertion, while LB-F and LB-R and RB-F and RB-R are two sets of primers used to identify random/extra insertion of T-DNA since they amplify T-DNA left border (LB) and right border (RB) repeats, respectively. Three exons and two introns of *ACO A* are marked by blue and gray rectangles, respectively. START and STOP indicate start and stop codon of *ACO A*, respectively. **b**, Confirmation of in-locus tagging of Aconitase A with GFP, and no extra random insertion of T-DNA in the selected strains using indicated primers. A bigger sized band (4974bp) will be amplified using ACOA 5’ UTR-F133 and ACOA 3’ UTR-R309 as primers if the in-locus tagging is successful, otherwise a smaller sized band (3034bp) or no band will be amplified. If extra/non-targeted insertion of T-DNA occurs, a 214 bp LB band and a 209 bp RB band will be amplified using indicated primers. Genomic DNA of strain #1 to #3 which shows Basta resistance was tested with 1 kb or 100bp DNA ladder included as a size reference. Wild type (WT) genomic DNA, water, or known genomic DNA that can function as negative (N) or positive (P) controls were also included in the test. **e**, Deleting *ATG24* through homologous recombination. *ATG24* Open reading frame starting from start codon (START) to stop codon (STOP) including 2 exons (blue rectangles) and 1 intron (gray rectangle) is replace by a sulfonylurea (specially chlorimuron ethyl) resistance cassette though homologous recombination as indicated. Primers used to verify *ATG24* deletion or confirm no random T-DNA insertion are also indicated. **f**, PCR based genotyping confirms successful *ATG24* deletion. An AcoA-GFP fusion band (711bp) was used as a loading control, and an 808 bp band will be amplified using primer ATG24-F and ATG24-R as indicated if *ATG24* deletion fails. **c**, Confirming mitochondrial localization of Aconitase A with TMRE staining. Projection images are shown. **d**, In-locus GFP tagging of the mitochondrial Aconitase does not affect its function as reflected by the similar ability to cause blast disease as compared with wild type (WT). Otherwise indicated, strain #3 was selected as a representative. **g**, AcoA-GFP localization with or without mitophagy at 7 to 8 hpi. Unless otherwise indicated, AcoA-GFP is shown as projections. Scale bars are 5 μm. Data shown in (**c**), (**d**), and (**g**) are representatives of at least two independent repeats with consistent results.

## Notes

https://doi.org/10.5281/zenodo.7943903

### Competing Interest Statement

The authors have declared no competing interest.

### Summary of Updates

Includes a detailed write-up for the Results section. A discussion section has been included too. Also revised the microscopy analysis for Figure 5 and 6.

